# COVID-19 PCR Test Performance for Samples Stored at Ambient Temperature

**DOI:** 10.1101/2020.06.15.153882

**Authors:** Nihat Bugra Agaoglu, Jale Yıldız, Ozlem Akgun Dogan, Gizem Alkurt, Betsi Kose, Yasemin Kendir Demirkol, Arzu Irvem, Levent Doğanay, Gizem Dinler Doganay

## Abstract

**Background:** The new type of Coronavirus infection had become a pandemic in a very short period since it was first seen in Wuhan. The outbreak had a negative impact on all health care systems throughout the world and overwhelmed the diagnostic laboratories as well. During the pandemic, handling patient specimens in accordance with the universal guidelines was troublesome as WHO, CDC and ECDC required cold chain compliance during transporting and storing the swap samples.

**Materials and methods:** In this study, we tested diagnostic performance of RT-PCR on 30 swab samples stored at ambient temperature and compared them with the samples stored at +4°C.

**Results:** Our results revealed that all the samples stored at ambient temperature remain PCR positive for at least five days. We did not see any false negativity.

**Conclusion:** In conclusion, we report that transferring and storing of nasopharyngeal/oropharyngeal samples at ambient temperature could be possible in the resource-limited conditions like pandemic.

## INTRODUCTION

Coronavirus disease 19 (COVID-19), which stems from a new type of coronavirus, severe acute respiratory syndrome coronavirus 2 (SARS-CoV-2), has become a universal pandemic since it first appeared in Wuhan, China in November 2019. Until the 10th of June, there were 7,127,753 confirmed cases, 407,159 deaths, and 108,918 new cases (WHO COVID-19 Dashboard) ^1^. With increasing number of infected people in a such short period of time, the pandemic has been an enormous burden to healthcare system including diagnostic laboratories. Real-time polymerase chain reaction (RT-PCR) is considered as the gold-standard confirmatory diagnostic laboratory test for CoVID-19 ^2^. Samples for this method are obtained from the upper and lower respiratory tract (oropharyngeal, nasopharyngeal swabs, sputum, lower respiratory tract aspirates, bronchoalveolar lavage, and nasopharyngeal wash/aspirate or nasal aspirate)^3^ and put into a transport medium. The World Health Organization (WHO), Centers for Disease Control and Prevention (CDC), European Centre for Disease Prevention and Control (ECDC) and several national health authorities announced several RT-PCR protocols and sample storage guidelines and all emphasized that accuracy of the RT-PCR tests mostly relies on proper specimen collection and storage ^4-6^. All guidelines agreed that samples should be transferred and kept in conditions maintaining cold chain. However, processing outrageous number of samples during the outbreak hampers keeping transport and storing conditions in line with the guidelines. Here, we aimed to test the RT-PCR diagnostic performance on nasopharyngeal/oropharyngeal swab samples stored at ambient temperature.

## MATERIAL and METHODS

### Sample Collection, Transportation, and Storage

This study was approved by the Umraniye Teaching and Research Hospital ethical committee Nasopharyngeal/oropharyngeal swabs were collected by trained personnel and transferred to GLAB-Corona ^7^ in Viral Transport Media (VTM, Innomed VTM001). Thirty samples tested positive with the quantification cycle (Ct) values between 12.41-23.12 were selected as study samples. 2 mL of VTM solution was added to the samples. Then these tubes were vortexes and aliquoted into two separate tubes (2 mL of solution for each tube) to be stored at refrigerator (+4°C) and room temperature (20-24°C), respectively. Additionally, five RT-PCR negative samples were stored under the same conditions as a control group. The room temperature and refrigerator temperature were monitored continuously and recorded using data loggers (Fig. 1).

**Fig 1.**
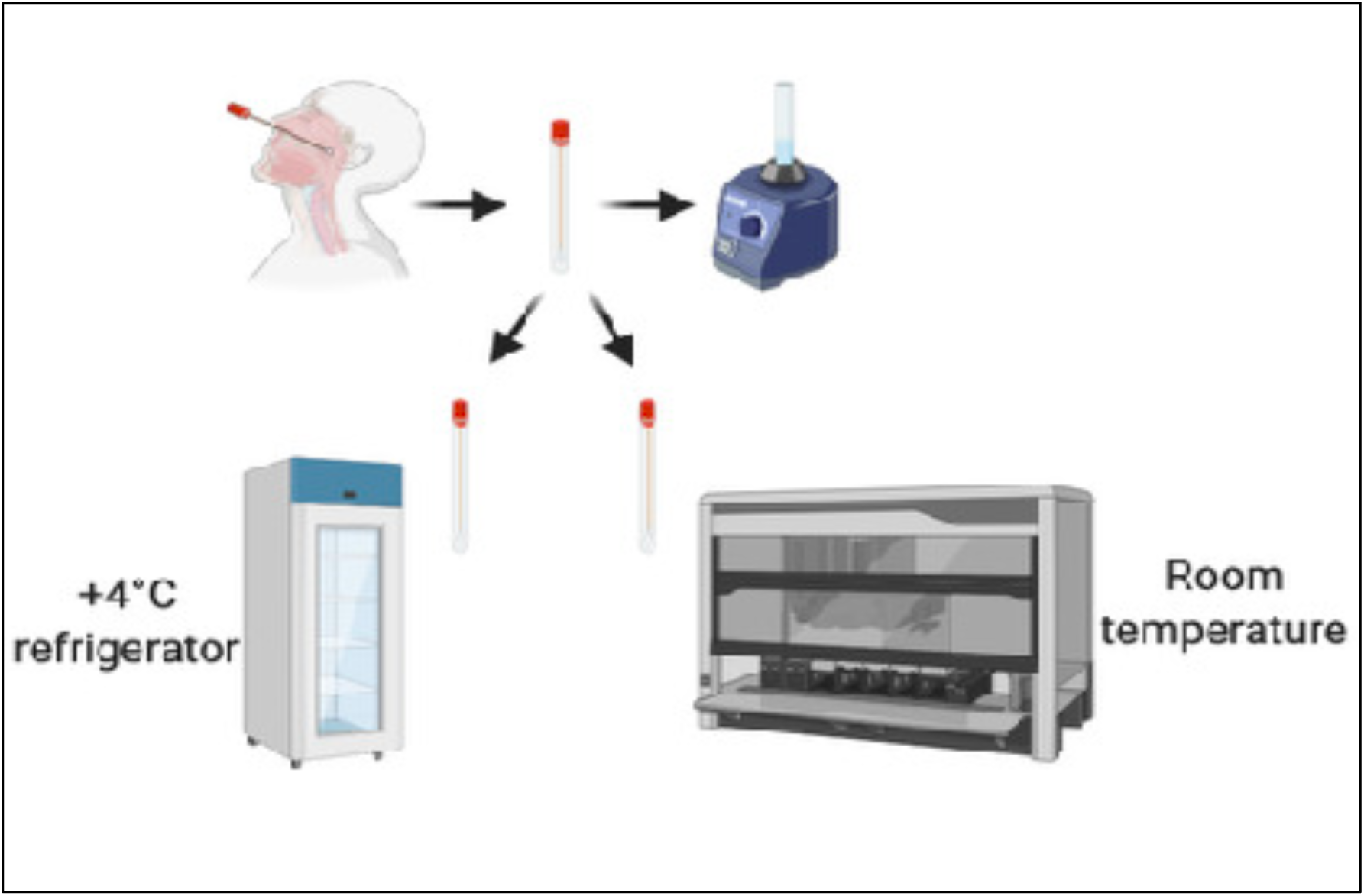
Sample storage workflow

### RT-PCR Tests

All samples (+4°C, room temperature, and controls) were tested every 24 hours for 9 days by RT-PCR with the SARS-CoV-2 detection kit (Coyote Bioscience Co., Ltd) according to the protocols provided by the manufacturer.

The primers of the commercial kit designed to target *ORF1ab* and *N gene* of SARS-CoV-2. All tests were done with Biorad CFX 96 platform. The cut off of Ct value was taken below 29 for the detection of positive results according to the suggestions of the manufacturer.

### Interpretation of RT-PCR Results

A positive result is reported if both the *ORFlab* and *N gene* were positive (Fig. 2). If both targets were negative, the result is negative. When only one target region is positive then the test was repeated.

**Fig 2.**
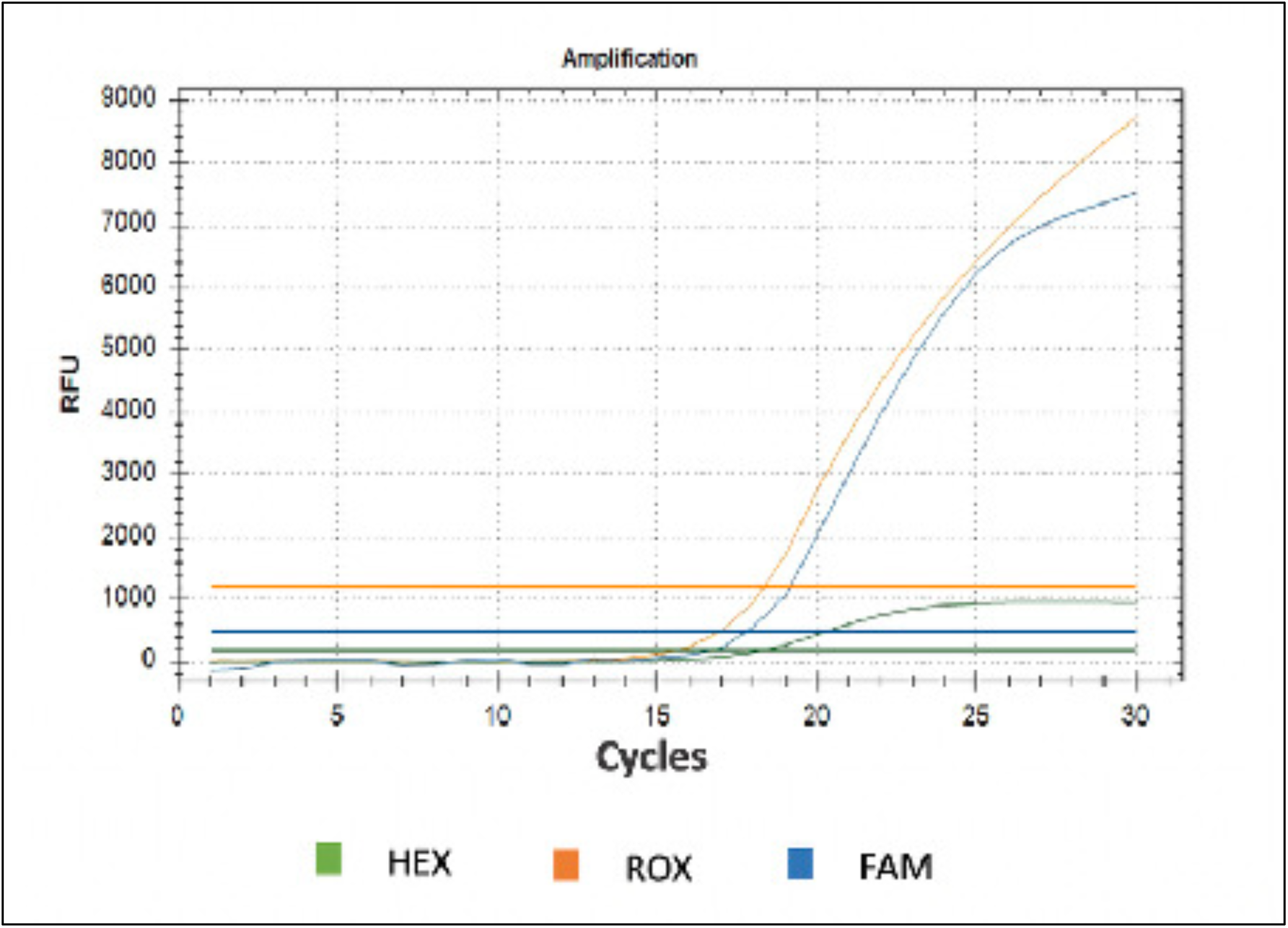
Amplification plot of a positive PCR result. To decide the positivity of the samples, FAM channel for *ORFlab* gene, ROX channel for *N gene*, and HEX channel for the internal RNase P gene of human control is evaluated. For a positive result the cut off for Ct is below 29.

## RESULTS

A total of 30 SARS-CoV-2 positive samples and 5 negative samples (all stored in +4°C and ambient temperature) were studied. All positive samples were run in PCR every day, and negative samples were studied on day 5 and day 9.

On day 6, 2 samples (in positive group) failed to run due to insufficient material. On day 7, 12 samples, on day 8, 17 and on the 9th day 28 samples had insufficient material for the PCR assay. Study is terminated at day 9 (Table 1).

**Table 1.** Daily PCR results of samples stored at +4°C and ambient temperature

|  | +4°C |  |  |  |  | Room Temperature |  |  |  |  |
| --- | --- | --- | --- | --- | --- | --- | --- | --- | --- | --- |
|  | Available<br>Sample<br>(n) | Positive<br>(n) | Negative<br>(n) | TR<br>(n) | SR<br>(n) | Available<br>Sample<br>(n) | Positive<br>(n) | Negative<br>(n) | TR<br>(n) | SR<br>(n) |
| **Day 0** | 30 | 30 | 0 | 0 | 0 | 30 | 30 | 0 | 0 | 0 |
| **Day1** | 30 | 30 | 0 | 3* | 0 | 30 | 30 | 0 | 4* | 0 |
| **Day2** | 30 | 29 | 1 | 1* | 0 | 30 | 30 | 0 | 1* | 0 |
| **Day3** | 30 | 29 | 1 | 3* | 0 | 30 | 30 | 0 | 0 | 0 |
| **Day4** | 30 | 30 | 0 | 0 | 0 | 30 | 30 | 0 | 0 | 0 |
| **Day5** | 30 | 29 | 1 | 2* | 0 | 30 | 30 | 0 | 0 | 0 |
| **Day6** | 28 | 27 | 0 | 0 | 1 | 30 | 30 | 0 | 0 | 0 |
| **Day7** | 22 | 22 | 0 | 0 | 0 | 24 | 24 | 0 | 0 | 0 |
| **Day8** | 15 | 15 | 0 | 0 | 0 | 23 | 23 | 0 | 0 | 0 |
| **Day9** | 5 | 3 | 2 | 0 | 0 | 12 | 12 | 0 | 0 | 0 |

\*All repeated tests resulted positive. TR: test repeat, SR: sample repeat

Five PCR negative samples were aliquoted and stored in refrigerator and ambient temperature as control samples. These controls tested on day 5 and day 9, all remained negative. The cycle values obtained are indicated in the tables (Table S1, Table S2, Table S3, Table S4).

PCR results of daily tested study samples are given in Table 1. On day 1, 3 samples in +4°C group and 4 samples in RT group resulted as “test repeat”. Ultimate PCR results for all these samples were positive. On day 2, 3 and 5, only one sample in +4°C group tested negative, interestingly this sample showed a positive result on day 4. After day 7 the available test material in some tubes remained insufficient for the PCR assay. Till day 7 test sensitivity in ambient temperature was 100%. We did not observe false negative in room temperature group even for the sample showing variation at +4°C.

Samples stored at +4°C showed instability in the test results by day 9 and also remaining samples in tubes were not sufficient for continuing PCR testing. On the other hand, samples kept at room temperature revealed higher consistency without showing any false negativity.

## DISCUSSION

Here we analyzed the RT-PCR diagnostic performance for SARS-CoV-2 on nasopharyngeal/oropharyngeal swab samples stored at ambient temperature. While many studies have investigated the effects of environmental factors such as temperature and humidity on virus survivability, less is known about the impact of temperature on SARS-CoV-2 RNA detection by RT-PCR ^8-10^. In a limited number of studies in this area related to other viruses (enterovirus, HSV-2, HHV-8, adenovirus, influenza), it has been reported that the storage of samples at ambient temperature did not affect the positive test results. Our results clearly show that oropharyngeal/nasopharyngeal samples kept at room temperature remain positive in SARS-CoV-2 RT-PCR studies for at least five days in accordance with these studies ^11, 12^.

CDC, WHO and ECDC recommend storing samples at 2-8°C for up to 3 and 5 days, respectively. Also, they all suggest storing at −70°C for samples that needs to be stored for more than 5 days or whose transfer will be delayed. However, this restrictive suggestion causes logistic and cost problems related to the transportation and storage of samples especially in pandemic periods when massive amounts of samples have to be analyzed. In addition, many laboratories, following the CDC and WHO recommendations, reject samples that cannot be delivered to them under the recommended transport conditions. This situation requires resampling, where both creates discomfort for the patient and causes delays in diagnosis.

It is obvious that keeping the number of RT-PCR tests as high as possible is the most crucial step in preventing the spread of infection. For this purpose, it is also very important to use existing infrastructure in an efficient way. In light of the data we obtained from our study, we suggest that in a resource-limited setting like pandemics, transferring and storing of nasopharyngeal/oropharyngeal samples at ambient temperature (20-24°C) should not be considered as inadequate, and these samples should not be rejected, can be analyzed and reported. Such a change in daily practice will result in considerable time and cost savings as well as a reduced number of sample rejection. When sample transportation or storage at cold chain conditions becomes a limiting factor for the pandemic laboratory, keeping samples at ambient temperature will enable much more testing.

The limitation of our study is the considered range of temperatures, between 20°C and 24°C. Temperatures higher than 24°C may be common in some tropical research and/or storage settings so the time period that samples remain positive may be variable. Also, the limited number of samples can be counted as another limitation of our study.

## ABBREVIATIONS

GLAB-Corona: COVID-19 Laboratory supported by the Health Institutes of Turkey (TUSEB) at the Umraniye Teaching and Research Hospital, Istanbul, Turkey.
RT-PCR: Real time polymerase chain amplification
Ct: Threshold Cycle
VTM: Viral transport medium

## ACKNOWLEDGMENTS

We thank Yesim Karaca, Aleyna Karaca, Irem Yaren Yar and Murat Kaya for the support and contributions for sample collection at the Genomic Laboratory, GLAB-Corona.

